# Comparative study of nanopore phenylalanine clamp variants reveals unique peptide biosensing and classification properties

**DOI:** 10.1101/2025.08.22.671566

**Authors:** Jennifer M. Colby, Bryan A. Krantz

## Abstract

The rapid, label-free, and low-cost detection of peptides is critical for the development of next-generation diagnostics, drug discovery, and environmental monitoring. Nanopore-based biosensing offers a promising platform to address this need by leveraging single-molecule analysis. In this study, we utilize protein engineering to create a series of novel peptide biosensors from the anthrax toxin protective antigen (PA) nanopore by targeting its central phenylalanine clamp constriction (residue F427), a key site known to interact dynamically with translocating molecules. This series of engineered variants were evaluated for their performance in both unsupervised clustering and supervised classification of a diverse set of seven guest-host peptides. Intriguingly, we found that the engineered variants exhibited a broad range of unique biosensing and classification properties. There was a notable divergence between the ability of the variants to intrinsically separate peptides (unsupervised clustering) and their performance in supervised classification tasks. Notably, PA F427A nanopores showed enhanced specificity for small molecular weight peptides that were challenging for WT nanopores to classify, achieving exceptionally high performance (accuracy of 0.93). These findings challenge the assumption that a single unmodified biosensor is sufficient for complex discrimination. Instead, our results highlight the potential for a more robust approach: leveraging the unique, complementary strengths of multiple sensor variants in an ensemble or multiplexed array. Such a system can achieve high and balanced performance across diverse peptide classes, representing a significant step forward in the development of sophisticated nanopore biosensors.

## Introduction

The precise identification and quantification of peptides and proteins are fundamental to a wide range of applications, from medical diagnostics to environmental monitoring. Peptide biomarkers hold immense promise for early detection of diseases, like heart disease, cancer, and infectious diseases (1–3). However, effectively analyzing complex and often dilute peptide mixtures demands analytical platforms with extraordinary sensitivity and specificity. While many conventional techniques struggle with these requirements, single-molecule nanopore biosensing offers a powerful, label-free alternative (4). This technology operates by detecting discrete changes in ionic current as individual molecules pass through a nanometer-scale pore, providing a real-time high-resolution readout of molecular properties. Beyond simple detection, this platform is a leading candidate for the ambitious goal of direct, high-throughput peptide and protein sequencing—an area of research that still lacks a widely applicable solution (5–7).

At the heart of nanopore sensing are biological protein channels, such as anthrax toxin PA, which provide a highly controlled and tunable environment for molecular interactions (8–13). These pores, when embedded in a membrane, can capture the dynamic, picoamp-scale current fluctuations, as translocating molecules interact with the nanopore’s internal features. The resulting ionic current signatures are rich in information, but their complexity presents a significant challenge, namely in extracting adequate analytical detail to enable reliable discrimination of diverse sets of peptide analytes. Critically, a single unmodified biosensor, regardless of its performance, may not possess the necessary discriminatory power to accurately classify the wide structural and chemical diversity found in real-world peptide mixtures. This suggests a need for a new paradigm in nanopore biosensor design.

The anthrax toxin PA channel has emerged as a premier biological nanopore model system for polypeptide analysis (14). As a natural protein translocase, PA is exceptionally robust and facilitates the processive, voltage- and/or proton gradient-driven transport of peptides (15, 16). It achieves this without the need for DNA tethers or other labels (17–20). PA uses multiple internal loops and clefts, called ‘peptide-clamp’ sites, such as the phenylalanine clamp (ϕ clamp), to interact with the translocating molecule (9, 12, 13, 18, 20, 21). These interactions generate complex, multi-state ionic current signatures with distinct kinetic and conductance characteristics (20), which contain the information necessary for both peptide classification (22) and, in principle, sequencing. The inherent structural modularity of the PA nanopore makes it an ideal platform for protein engineering to fine-tune its biosensing properties.

Analyzing the voluminous and often noisy data from these multi-state systems necessitates advanced computational methods. Machine learning (ML) and deep learning are particularly well-suited for this task, as they can identify subtle, high-dimensional patterns in translocation kinetics that are inaccessible to traditional analysis (22–28). However, the full potential of these computational tools has yet to be realized, especially when dealing with the challenge of analyzing complex peptide mixtures.

In this work, we hypothesize here that a single unmodified sensor, optimized for certain types of peptides, may not perform as successfully across a wider array of different analytes. To address this, we are leveraging the PA nanopore as a model system, employing protein engineering of the ϕ-clamp site to generate a series of novel biosensors. We then use advanced ML methods to characterize the unique classification strengths of each variant. Our results demonstrate that an ensemble or multiplexed array of these engineered biosensors, with their complementary detection characteristics, can achieve a well-balanced performance, surmounting the performance of the wild-type (WT) pore. This ensemble approach represents a significant step forward for protein nanopore-based analysis in biotechnology, paving the way for more accurate peptide and protein sequencing capabilities.

## Results

### Nanopore variant study design

Our study leveraged the well-characterized anthrax toxin PA nanopore as a single-molecule peptide biosensor. The PA nanopore’s architecture includes a key narrow constriction site, known as the ϕ clamp, which is formed by a radially symmetric arrangement of phenylalanine residues (F427) (13). This site is positioned to make close contact with translocating proteins, effectively ‘clamping’ them and thereby promoting productive protein unfolding and translocation. Moreover, the interaction leads to full and partial blockade of the channel to ion flow in a highly dynamic and peptide-specific manner (20, 22). Our central hypothesis was that tuning the identity of the residue at this F427 position would alter the nanopore’s selectivity and, consequently, its capabilities as a peptide classifier. To test this, we engineered two specific variants (i) F427A (replacing Phe with the much smaller Ala) and (ii) F427Y (replacing the site with the slightly bulkier, yet more hydrophilic Tyr). We then compared their classification performance to the WT PA nanopore.

To evaluate our hypothesis, we used a previously characterized set of seven 10-residue guest-host peptides of the sequence KKKKKXXSXX (where X is the guest residue) (17, 20, 22). The guest residues included non-aromatic side chains (Ala, Leu, Thr) and aromatic side chains (Phe, Trp, Tyr). A final variant, guest-host TrpDL, featured an alternating pattern of d- and ʟ-stereochemistry to probe how peptide backbone dynamics influence the translocation signal. Prior work has demonstrated that these peptides can be efficiently classified by the WT PA nanopore using ML with ∼0.90 accuracy (22), and we used these results as a benchmark for the engineered F427A and F427Y nanopores.

### Data collection and preprocessing

In previous work, we collected WT PA nanopore translocation event streams for the guest-host peptide series. These experiments were performed under a constant driving force of 70 mV (cis positive), a condition known to strongly favor productive translocation events and produce robust current signals (22). We similarly collected single-channel peptide translocation recordings for the F427A and F427Y nanopore variants under the same 70 mV driving force, with data streams extending up to 30 minutes per peptide. The raw data from these experiments consisted of complex ionic current signals **(Fig. 1)**. To prepare this data for ML, we developed a preprocessing pipeline. First, using a K-Means clustering algorithm, the raw event streams were labeled according to their corresponding conductance states. While rare intermediate states were occasionally observed, our analysis identified four robust and well-populated states that consistently described most of the current signal. These states are enumerated 0 through 3, corresponding to a fully blocked (state 0), partially blocked (states 1 and 2), and fully open (state 3). This state labeling procedure was benchmarked against expert electrophysiology software, CLAMPFIT, and deemed to be highly consistent yet computationally much more efficient. These state-labeled records were then carefully segmented into individual translocation events. We settled on using only events greater than or equal to 15 ms in length in all our subsequent analyses to capture the most data while excluding lower-information short duration spiky events. Finally, a set of biophysical features, such as dwell time, current blockade level, and signal variance, were computed to describe each individual event (see Materials and Methods for a complete list of features). These comprehensive feature sets, representing the unique biophysical signature of each peptide translocation event, were then used as the training data for our ML classification models.

**Fig. 1.**
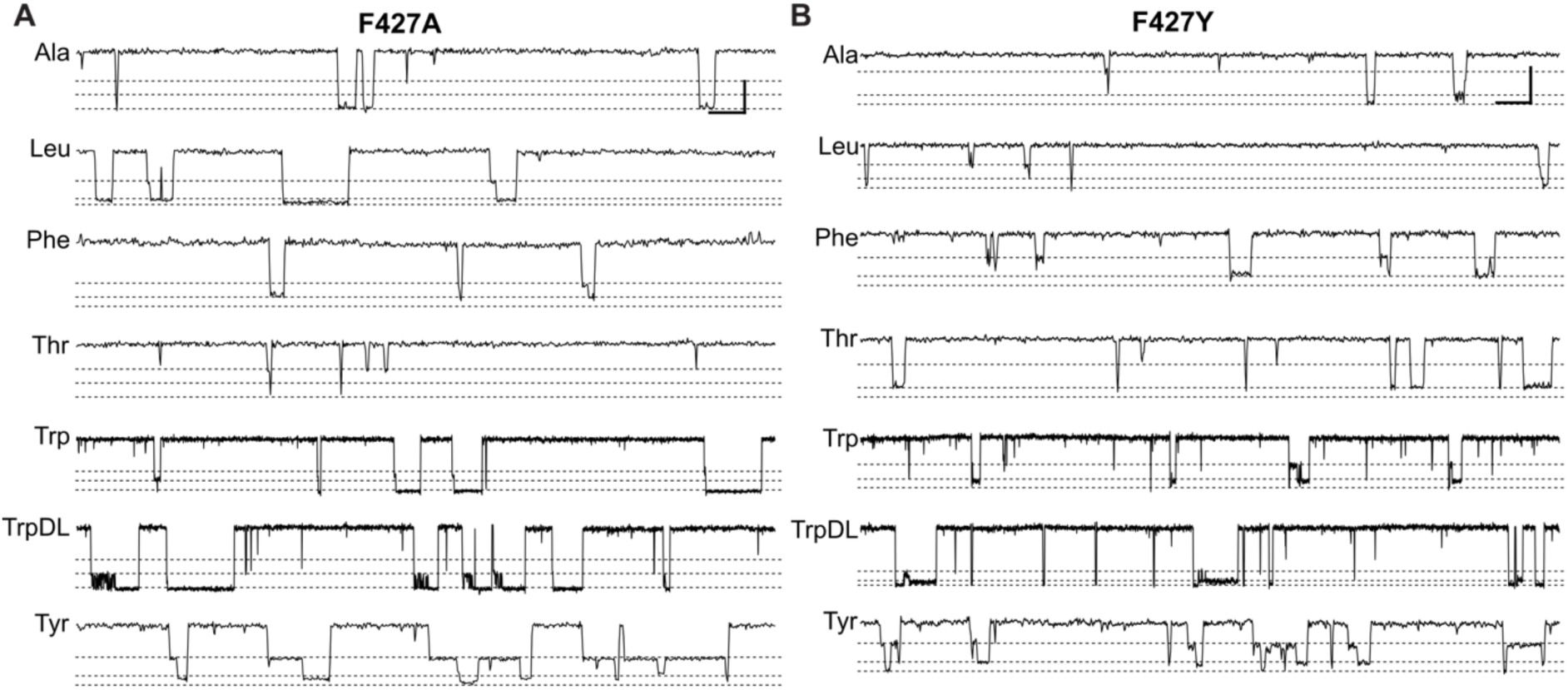
Translocation event streams for F427A and F427Y nanopores. Representative recordings of guest-host peptide translocations carried out at 70 mV (cis positive) in symmetric succinate buffer, 100 mM KCl, pH 5.6 for **(A)** F427A and **(B)** F427Y nanopores. To the left are the standard three letter name for the guest residue. These nanopore-peptide systems populate multiple discrete partially or fully blocked intermediates (approximate locations indicated by dashed lines). From bottom to top of each record: fully blocked (state 0), partially blocked intermediates (state 1 and state 2), and fully open baseline (state 3). Scalebar at the upper right of each panel denotes 4 pA by 100 ms for guest-host Ala, Leu, Phe, Thr, and Tyr peptides. For guest-host Trp and TrpDL peptides, it is 4 pA by 500 ms to show their characteristically longer events. Note well that F427A nanopores conduct ∼30% more than F427Y channels (13).

### Unsupervised clustering analysis

To assess the intrinsic separability of the peptide features extracted from each nanopore variant, we performed an unsupervised clustering analysis using Uniform Manifold Approximation and Projection (UMAP) **(Fig. 2)**. UMAP is a dimensionality reduction technique that is well-suited for visualizing the high-dimensional data, revealing the underlying structure and grouping of feature sets. To quantitatively evaluate the quality of the clustering, we calculated the Adjusted Rand Index (ARI) and Normalized Mutual Information (NMI) for each nanopore, comparing the UMAP-derived clusters against the ground truth peptide labels. In the analysis, using a 15 ms minimum event duration filter, WT PA nanopore had an ARI of 0.0648 and NMI of 0.0864; PA F427A had an ARI of 0.0536 and NMI of 0.0775; and PA F427Y had an ARI of 0.0947 and NMI of 0.1424. A higher ARI and NMI value indicates that the feature set of a given nanopore more effectively clusters the peptides into groups that align with their true identities. Visual UMAP plots for the F427Y variant corroborated these metrics, showing less scatter and cleaner clusters compared to the F427A variant **(Fig. 2)**. Interestingly, these results suggest that the features from the PA F427Y nanopore, which will be shown to have lower performance in supervised classification tests, demonstrate superior inherent clustering capabilities. This highlights a critical distinction between unsupervised and supervised learning tasks. While the features of the F427Y nanopore may create tight, well-defined clusters, these clusters may be positioned too closely to one another in the feature space, making it difficult for a supervised classifier to establish a clear decision boundary. Conversely, a nanopore with less tightly clustered data (like WT or F427A) may still have greater cluster separation, which is a more favorable condition for a supervised model to learn and generalize from. This observation underscores the fact that the ultimate metric of success in this study is not the inherent separability of the data, but the performance of the classification models built upon it. The following section will therefore focus on the supervised learning performance, where the true discriminative power of each nanopore variant is directly measured.

**Fig. 2.**
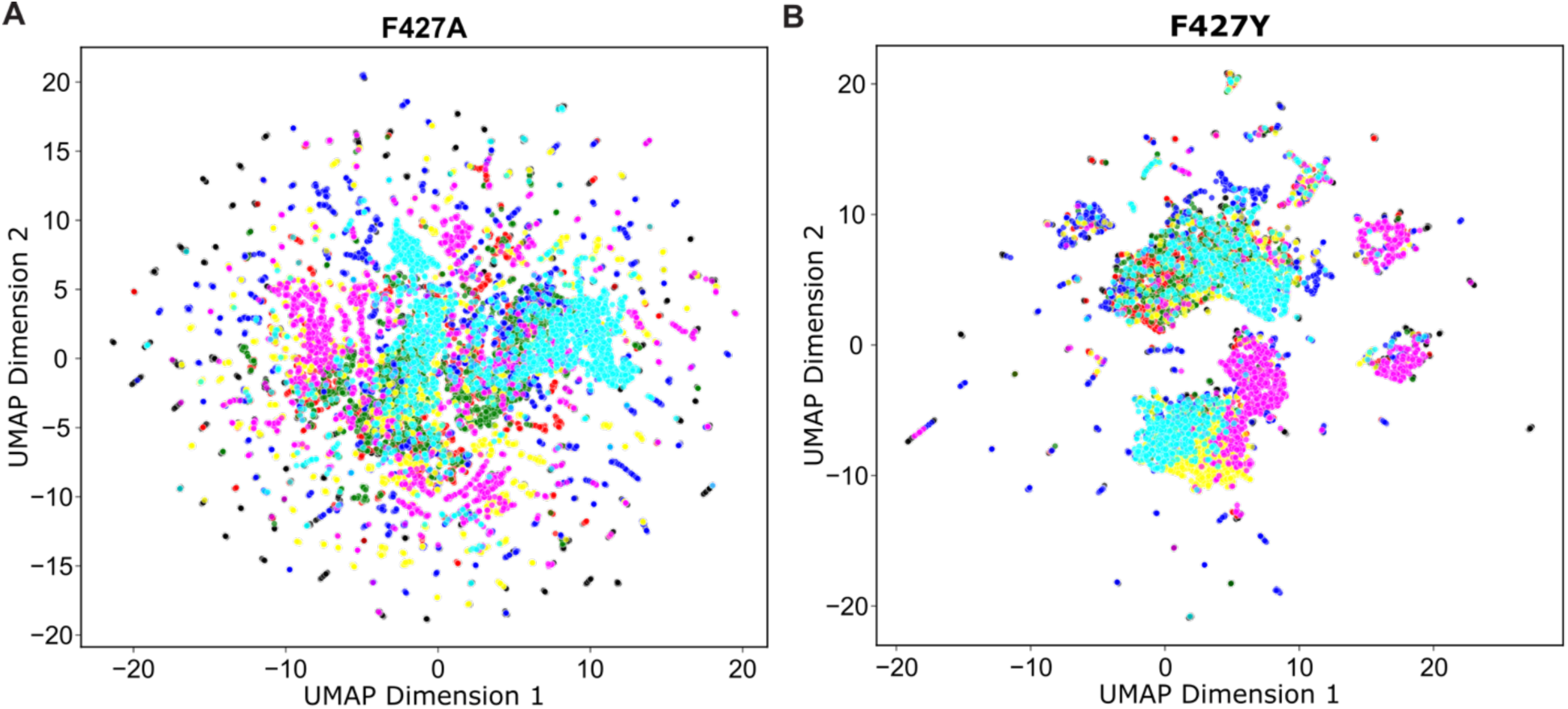
Unsupervised clustering analysis of feature set describing translocation events from mutant nanopores. UMAP clustering analysis of event-level features. Data points, representing individual translocation events, are colored by guest-host peptide identity: Ala (black), Leu (red), Phe (green), Thr (blue), Trp (yellow), TrpDL (magenta), and Tyr (cyan). **(A)** UMAP embedding of F427A nanopore with ARI of 0.0536 and NMI of 0.0775. **(B)** UMAP embedding of F427Y nanopore events with ARI of 0.0947 and NMI of 0.1424. Events in either panel were filtered at a minimum event duration of 15 ms, and UMAP settings were n_neighbors = 15 and min_dist = 0.5.

### Initial single-stage classification performance of nanopore variants

To establish a performance baseline, we first trained individual single-stage eXtreme Gradient Boosting (XGBoost) classifiers for each nanopore variant: WT, F427A, and F427Y **(Table S1)**. This preliminary analysis revealed distinct performance profiles for each nanopore, suggesting a potential for synergistic gains through an ensemble approach. The WT nanopore achieved a mean macro-averaged F1-score of 0.8652 (±0.0087) (N=5), while the F427A variant demonstrated superior overall performance with a mean F1-score of 0.8869 (±0.0089). In contrast, the F427Y nanopore performed the lowest, with a mean F1-score of 0.7829 (±0.0088).

A detailed, class-by-class analysis of the individual F1-scores revealed the specific strengths of each nanopore **(Fig. 3)**. The F427A nanopore demonstrated a marked increase in predictive power for guest-host peptides containing small non-aromatic side chains. Specifically, its F1-scores for guest-host Ala (0.95) and guest-host Thr (0.94) were substantially higher than those achieved by the WT nanopore (0.80 and 0.74, respectively) **(Table S1)**. Conversely, the WT nanopore showed a slight advantage in classifying some aromatic guest-host peptides, such as guest-host TrpDL, with an F1-score of 0.96 compared to 0.91 for F427A. The WT nanopore also performed well on guest-host Phe, Trp and Tyr. Confusion matrices similarly show the high degree of guest-host Ala and Thr confusion for the WT nanopore relative to the F427A version **(Fig. S1)**. These complementary strengths—F427A excelling with small non-aromatic side chains and WT performing somewhat better on aromatic ones—motivated the design of a more sophisticated, multistage ensemble classifier to leverage the unique information from each nanopore variant.

**Fig. 3.**
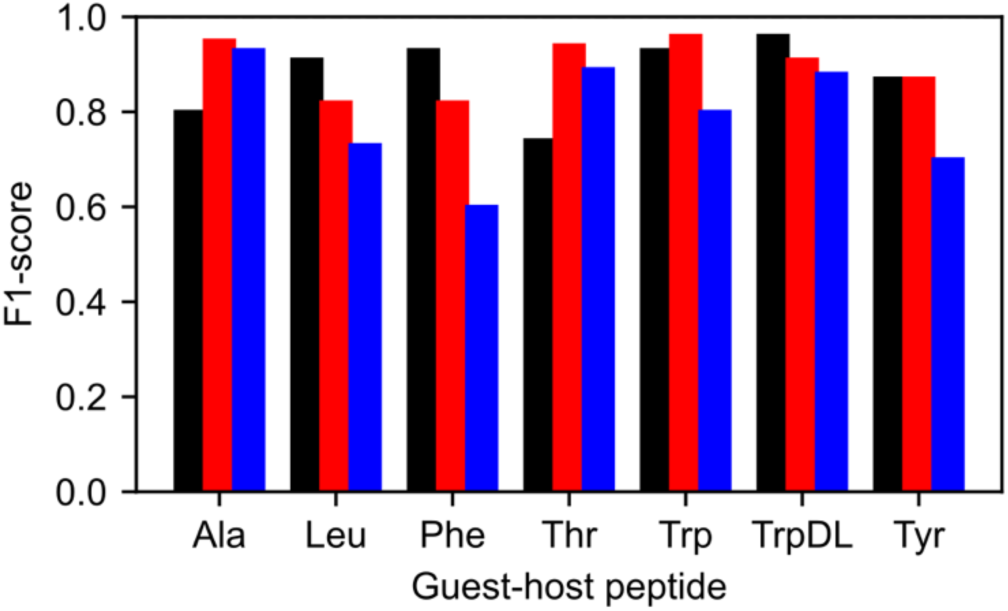
Per-peptide class F1-scores of individual nanopore ϕ-clamp variants. Per-class F1-scores for single-stage XGBoost classification for WT (black), F427A (red), and F427Y (blue) nanopores. Plotted values represent the best performing evaluation out of 5 replicate trainings.

### Multistage ensemble classifier architecture

To effectively maximize the distinct single-molecule information from each nanopore variant, we designed a multistage ensemble classifier based on a probabilistic blending approach **(Fig. 4A)**. In the first stage, individual XGBoost classifiers were trained for each nanopore variant (WT, F427A, or F427Y). These models were configured to output a probabilistic vector (softmax scores) indicating the confidence of an event being from an aromatic peptide class. In the second stage, a separate XGBoost model was trained on the augmented features, which now included the Stage 1 probabilities, to predict peptide-specific probabilities for both aromatic and non-aromatic peptides. Finally, a meta-classifier took as input the combined feature set from the raw data and the probabilistic outputs from both stages 1 and 2 to produce the final peptide predictions. For instance, in a two-nanopore combination (e.g., WT/F427A), the meta-classifier’s input vector was a blend of all softmax scores from both nanopores from both stages. This architecture allowed the meta-classifier to optimally blend the confidence scores from the different nanopore classifiers, effectively identifying and amplifying the unique and complementary information provided by each variant to improve overall classification accuracy.

**Fig. 4.**
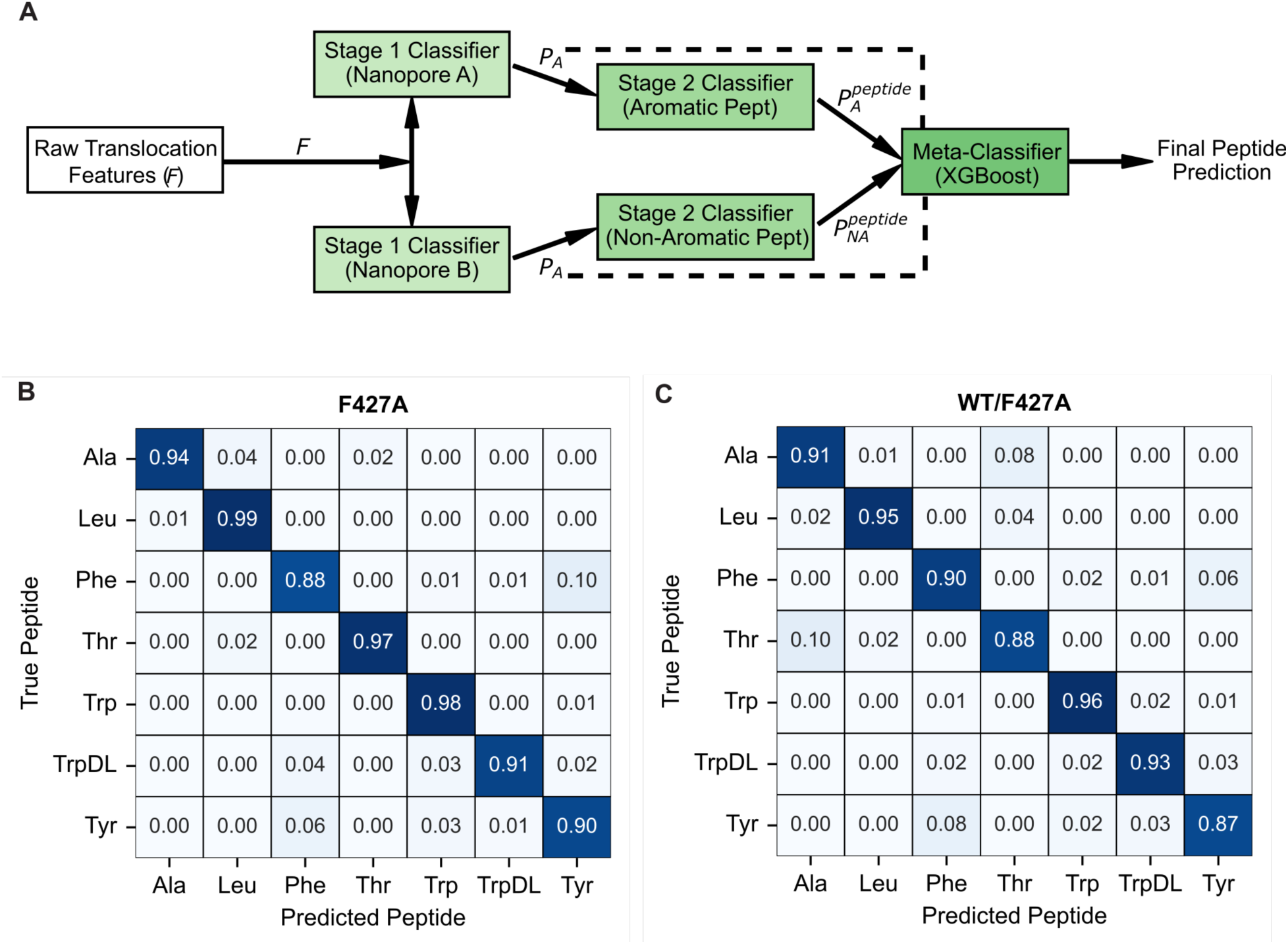
Multistage ensemble probabilistic blending classifier. **(A)** Block diagram of multistage ensemble probabilistic blending classifier. The model processes raw translocation features (*F*) through a two-stage classification and blending pipeline. Stage 1 classifiers, one for each nanopore variant, predict the aromatic class probability (*P_A_*) for a given translocation event. These probabilities are then used to augment the original features (*F*) for Stage 2 classification. The Stage 2 classifiers predict peptide-specific probabilities for both aromatic (*P_A_^peptide^*) and non-aromatic (*P_NA_^peptide^*) peptides. Finally, a Meta-Classifier blends the probabilities from both Stage 1 (dashed line connections) and Stage 2 to produce the final peptide prediction. Normalized confusion matrices from most successful trainings of multistage ensemble probabilistic blending model for **(B)** a single F427A nanopore and **(C)** a WT/F427A ensemble. Rows represent true peptide labels, and columns represent predicted labels. Values indicate the proportion of events from a given true class that were predicted as each class.

### Classification performance of single and ensemble nanopores

We evaluated using the multistage model the performance of each single nanopore and all possible multiplexed nanopore combinations using 5 replicates to calculate the mean and standard deviation for key metrics **(Table 1)**. Of the individual nanopore classifiers, the F427A variant demonstrated the highest performance, achieving a mean macro-averaged F1-score of 0.9322 (±0.0042) (N=5). The WT nanopore followed with a mean F1-score of 0.8870 (±0.0094), while the F427Y nanopore performed the lowest, with a mean F1-score of 0.8367 (±0.0047). The performance of the nanopore combinations highlighted the benefits of our ensemble approach. The two-nanopore combination of WT/F427A emerged as the top-performing ensemble, achieving a mean macro-averaged F1-score of 0.9112 (±0.0018). This result is a notable improvement over the WT nanopore alone, demonstrating that the ensemble method effectively utilizes the combined signal. Confusion matrices for F427A and the WT/F427A ensemble **(Fig. 4B,C)** reveal more highly balanced classification with less confusion than the unmodified WT nanopore using a single-stage classifier **(Fig. S1)**. Moreover, examining the per-class F1-scores revealed that the WT/F427A combination yielded more balanced classification than the unmodified WT nanopore **(Table 1)**. Interestingly, the WT/F427A combination outperformed the three-nanopore ensemble (WT/F427A/F427Y), which achieved a mean F1-score of 0.8847 (±0.0032). Furthermore, the other combinations (WT/F427Y and F427A/F427Y) consistently showed lower performance than the best WT/F427A ensemble.

**Table 1.** Multistage ensemble probabilistic blending classifier performance^1^.

| Nanopore(s) | Overall accuracy<sup>2</sup> | Macro-averaged F1-score<sup>2</sup> | F1-score per guest-host peptide class<sup>3</sup> |  |  |  |  |  |  |
| --- | --- | --- | --- | --- | --- | --- | --- | --- | --- |
|  |  |  | Ala | Leu | Phe | Thr | Trp | TrpDL | Tyr |
| WT | 0.8874<br>(±0.0094) | 0.8870<br>(±0.0094) | 0.81 | 0.95 | 0.93 | 0.77 | 0.95 | 0.98 | 0.91 |
| F427A | 0.9323<br>(±0.0041) | 0.9322<br>(±0.0042) | 0.96 | 0.96 | 0.89 | 0.97 | 0.96 | 0.94 | 0.89 |
| F427Y | 0.8365<br>(±0.0042) | 0.8367<br>(±0.0047) | 0.97 | 0.92 | 0.67 | 0.94 | 0.82 | 0.86 | 0.72 |
| WT/F427A | 0.9112<br>(±0.0019) | 0.9112<br>(±0.0018) | 0.90 | 0.96 | 0.90 | 0.88 | 0.95 | 0.93 | 0.88 |
| WT/F427Y | 0.8632<br>(±0.0060) | 0.8632<br>(±0.0062) | 0.86 | 0.93 | 0.83 | 0.83 | 0.89 | 0.92 | 0.84 |
| F427A/F427Y | 0.8909<br>(±0.0061) | 0.8909<br>(±0.0061) | 0.97 | 0.94 | 0.81 | 0.96 | 0.90 | 0.91 | 0.82 |
| WT/F427A/F427Y | 0.8848<br>(±0.0033) | 0.8847<br>(±0.0032) | 0.89 | 0.94 | 0.85 | 0.88 | 0.91 | 0.91 | 0.86 |

<sup>1</sup>Performance based on test set evaluation metrics of multistage ensemble probabilistic classification model.
<sup>2</sup>Overall accuracy and macro-averaged F1-score values are means and std. dev. (N=5).
<sup>3</sup>Best set of per-class F1-scores out of N=5 replicates.

## Discussion

### Role of the ϕ clamp in protein and peptide translocation

The anthrax toxin PA nanopore’s constriction site, called the ϕ clamp, is a critical determinant of its biophysical properties and its ability to sense translocating molecules (13, 18, 20). This site, formed by a ring of phenylalanine residues (F427 in the WT oligomer), is the narrowest point of the pore with a luminal diameter of 6 Å. Its hydrophobic and aromatic character facilitates specific π−π stacking, π-cation, π-dipole, and hydrophobic interactions with passing substrates. While this constriction is essential for sensing, it also may present a significant energetic barrier for the unfolding and translocation of larger peptides and proteins. Our work with the F427A mutant demonstrates this nuance: while this mutant can still readily translocate peptides (18, 20), its reduced functionality can impede the passage of larger proteins at low driving forces (13). It is important to note, however, that at higher applied voltages, the F427A channel can still translocate large proteins, highlighting the complex interplay between molecular size, applied force, and the biophysical properties of the nanopore constriction. These observations set the stage for our central hypothesis: that engineered F427X variants could be uniquely tuned to create highly effective peptide biosensors.

### Correlating nanopore chemistry to peptide classification performance

Our findings reveal a clear correlation between the specific chemistry of the ϕ-clamp and the ability of the PA nanopore to distinguish between different peptides. The raw confusion matrices from the single-nanopore XGBoost models serve as a key example of this. The WT PA nanopore, while a capable sensor, showed poor performance and high confusion for the guest-host Ala and guest-host Thr peptides, which lack bulk and aromaticity (22) **(Table S1) (Fig. S1)**. In contrast, the engineered F427A and F427Y variants demonstrated distinct capabilities, including marked improvement in guest-host Ala and guest-host Thr predictions **(Table S1) (Fig. S1)**. By altering the specific side chain at the constriction, we were able to change the nature of the molecular interactions, which in turn resulted in unique current signals. This highlights how an engineered PA channel can exhibit a heightened ability to detect specific peptide chemistries that may be poorly resolved by the unmodified WT channel, providing more nuanced and effective classification.

### Beyond WT: value of engineered nanopore biosensors

Our work underscores the significant value of empirically testing engineered nanopore variants rather than relying solely on the WT PA nanopore for biosensor development. Counter-intuitively, we found that the F427A variant, which is known to translocate large proteins less efficiently than its WT counterpart, serves as a more valuable biosensor for our model peptides. This highlights a crucial design principle: a biosensor’s performance is not necessarily tied to its ability to translocate all molecules with high efficiency. Instead, its value lies in its capacity to generate unique, distinguishable signals for the specific analytes of interest. In the long run, our results suggest that a multiplexed array or ensemble of PA nanopores, each with a uniquely engineered constriction site, may be the most advantageous approach for creating robust biosensors that can cover a fuller breadth of peptide chemistries found in the real world.

### Future implications for nanopore-based peptide sequencing

The insights gained from this study have important implications for the future of nanopore-based peptide sequencing. Our results suggest that smaller residues at position 427, such as F427A, may be predicted to provide greater detail in sequencing applications. A smaller side chain there, making a less restrictive pore, might produce a more resolved signal as each amino acid passes. While here we have taken advantage of ML models for high performance biosensing, arguably sequencing may be more effectively tackled with high-throughput data collection, where deep learning models can be used to learn patterns and map current sequences to peptide sequences. Finally, the PA nanopore already possesses several qualities advantageous for sequencing development: it is already an electrophysiologically proven label-free protein (13, 15, 16) and peptide (17–20, 22) translocase; it senses at nanomolar peptide concentrations (18–20, 22); it is a capable biosensor platform (22); and it can be engineered in its loop and clamp sites for unique properties (12, 13, 21, 29), a key step in many sequencing applications.

### Future directions

The success of nanopore engineering here combined with the computational power of efficient multistage ensemble classifiers lays the groundwork for further development. Clearly, work can be done to expand the peptide library, including not only adding other side-chain chemistries in the guest-host series (exploring charge, hydrophilicity or hydrophobicity, and size) but also incorporating more natural sequences from real proteins. How generalizable are the conductance state intermediates observed in other peptide systems? Which nanopore variants are best suited to cover the most sequence space? This broadened approach will enable a more comprehensive understanding of our sensing capabilities for F427X engineered nanopores. Our multistage model reports feature importance, which may be used as a guide in the rational design of new nanopore mutants, creating a potentially iterative loop between computational prediction and experimental validation. Future work should also apply this multistage ensemble model to a wider range of nanopore variants or even to different nanopore systems to assess its generalizability for enhanced biosensing. Of course, real-world application of nanopore biosensors should involve research into resolving complex mixtures of peptides, including those found in biological fluids. Our work on individual event level predictions (22, 28) is a key step in detecting peptides from isolated events within mixed samples, but these computational approaches need to be met with analysis of real-world biological specimens.

## Materials and Methods

### Nanopore and peptides

Monomeric 83-kDa PA (PA_83_) preprotein mutants, F427A and F427Y, and their homoheptameric prepore oligomers (PA_7_) were produced as described (13, 30). PA_83_ mutants were overexpressed in *Escherichia coli* BL21(DE3), using a pET22b plasmid, which directs expression to the periplasm. Cell cultures were grown at 37 °C in a custom 5 L fermentor using ECPM1 growth media (31), which was supplemented with carbenicillin (50 mg/L). Once reaching an OD_600_ of 3-10, the cultures were then induced with 1 mM isopropyl β-d-thiogalactopyranoside for ∼3 h at 30 °C. PA_83_ was released from the periplasm by resuspending pelleted cells on ice using a wire whisk with 1 L of hypertonic sucrose buffer (20% sucrose, 20 mM Tris-Cl, 0.5 mM EDTA, pH 8) followed by osmotic shock of centrifuged/pelleted cells using a wire whisk in 1 L of hypotonic solution (5 mM MgCl_2_). Released PA_83_ monomer, isolated after centrifugation to remove cellular debris, was purified on Q-Sepharose anion-exchange chromatography in 20 mM Tris-Cl, pH 8.0 by binding and then eluting with a linear salt gradient using 20 mM Tris-Cl, pH 8.0 with 1 M NaCl.

To make PA_7_ prepore oligomers of either mutation, purified PA_83_ at a concentration of 1 mg/ml was treated with trypsin (1:1000 wt/wt trypsin:PA) for 30 min at room temperature to form nicked PA. Trypsin was subsequently inhibited with soybean trypsin inhibitor at 1:100 dilution (wt/wt soybean trypsin inhibitor:PA). Nicked PA was applied to Q-Sepharose to then isolate the PA_7_; oligomer was bound to the column in 20 mM Tris-chloride, pH 8.0 and eluted by a linear salt gradient using 20 mM Tris-Cl, 1 M NaCl, pH 8.0. PA_7_ was concentrated and frozen in small aliquots to maintain reproducible nanopore insertion activity in planar bilayer experiments.

Ten-residue guest-host peptides of the general sequence, KKKKKXXSXX, where X = A, L, F, T, W, and Y, were synthesized with standard ʟ amino acids (17, 20) (Elim Biopharmaceuticals). One stereochemical variant of X = W (called TrpDL) was produced, where instead of synthesizing the peptide with uniform ʟ amino acids, an alternating pattern of d and ʟ amino acids was used (20).

### Single-channel electrophysiology

Planar lipid bilayer currents were recorded using an Axopatch 200B amplifier interfaced by a Digidata 1440A acquisition system (Molecular Devices) (20, 30, 32). Membranes were formed by painting across a 50-μm aperture of a 1-mL white Delrin cup with 3% (wt/vol) 1,2-diphytanoyl-*sn*-glycero-3-phosphocholine (Avanti Polar Lipids) in *n*-decane. The *cis* (side to which the PA_7_ is added) and *trans* chambers were bathed in symmetric single-channel buffer (SCB: 100 mM KCl, 1 mM EDTA, 10 mM succinic acid, pH 5.60). Recordings were acquired at 500-600 Hz using PCLAMP10. The applied voltage is defined as Δψ = ψ_cis_ - ψ_trans_ (where ψ_trans_ is 0 mV).

Single-channel recordings of the guest-host peptide translocations via the PA nanopore were carried out as described (20) with some slight differences. A single PA channel was inserted into a painted bilayer at a Δψ of 20-30 mV by adding ∼2 pM of PA_7_ (freshly diluted from a 2-μM stock) to the *cis* side of the membrane. The prepore oligomer converts to the nanopore state by inserting into the membrane in an oriented manner. Once a single channel inserted, the *cis* chamber was perfused by fresh SCB to remove excess uninserted PA_7_. Then the desired peptide analyte was added to the *cis* chamber at 5 to 20 nM. Translocation data were acquired by stepping the applied Δψ to a higher positive value and collecting recordings of the translocation event stream for up to thirty minutes.

Minor processing as well as conductance state labeling of the raw single-channel event stream recordings was subsequently performed. Rare transient out-of-range current spikes, insertion of second channels, and inactivated channels were removed by a ‘force values’ routine in CLAMPFIT. Translocation recordings acquired at 500 or 600 Hz were downsampled to 400 Hz by decimation in Python using the scipy.signal library. 400 Hz was chosen to be consistent with prior WT PA datasets (22), maintaining a consistent time step for ML peptide classification models. Four discrete conductance states were then detected in these recordings using K-Means clustering, where baseline current drift was corrected by applying a moving window average offset (4000 time point window). While rare additional states were noted, the data were labeled for the four dominant states for either the F427A or F427Y mutants. By convention, the fully blocked peptide-bound state was state 0, the intermediate closest to the fully blocked state was state 1, the intermediate closest to the open state was state 2, and the open state was state 3. Ultimately, the state labeling produced a three-column CSV file of the stream with columns ‘Time’, ‘Current’, and ‘State’. >90% of time points were labeled consistently when comparing our K-Means algorithm and the more traditional CLAMPFIT. All labeled CSV stream files for the seven peptides were entered into a local annotated peptide database to aid in *in situ* loading/preprocessing for each tested ML model.

### Hardware and software used for ML

Anaconda was used to create a Python 3.10.16 environment, where XGBoost (3.0.0) (33) and other standard modules were installed. The hardware used in preprocessing and the ML peptide classifications was a 2025 MacBook Pro with M4 Apple Silicon and 24 GB of RAM. All source code is available at GitHub (https://github.com/bakrantz/Pept-Class).

### Preprocessing, event segmentation, and feature extraction

Raw state-labeled event streams were segmented into translocation events, enabling feature extraction, as described (22). The minimum event duration, which served as an effective filter for excluding very short-duration events, was set to 15 ms to optimally capture the most information-rich data. This filtering parameter was previously systematically varied in the range of 5 to 20 ms (22). Each segmented event was defined as initiating when the current changed from the fully open state (state 3, corresponding to baseline current) to any peptide-bound state (state 0, 1, or 2) and terminating when the current returned to the open state. From these segmented events, both raw current sequences and corresponding state sequences were extracted. A comprehensive set of event-level features was then computed from these two different sequences using a custom segmentation core. This core maintains a generalizable framework to process peptide translocation events from systems exhibiting diverse mechanisms and an arbitrary number of states. These features were initially grouped into scalar, vector, and matrix data structures, with values in vector and matrix features being state or transition enumerated. Scalar features included: Shannon entropy of state sequence, event duration, number of transitions, time of the first transition, total number of states visited during the event, skewness of the current sequence, and kurtosis of the current sequence. Vector features included: observed conductance state Boolean, observed conductance levels, probability of residing in each state, and longest dwell time in each state. Matrix features included: average dwell time for specific state-to-state transitions, variance of dwell time for transitions, and ratio of probabilities between states. For downstream ML classification, all matrix features were flattened into one-dimensional arrays and appended with the vector and scalar features to form a single feature vector for each translocation event. These flattened descriptive key names for the features were generated to maintain traceability in subsequent applications. All processed event sequences, their flattened features, and associated feature key names were saved as a Python pickle object for efficient storage and cached retrieval. A local peptide events database was employed to track these preprocessed cached pickle files for efficiency.

### Initial single nanopore ML peptide classification

ML-based classification of peptide translocation events from single individual nanopore variants was performed using the gradient boosting framework, XGBoost (33), which was implemented as described (22). For this, peptide event data, previously extracted and characterized into event-level features, were loaded from a local database. To ensure balanced class representation, all peptide classes were downsampled to match the class with the minimum number of events. The comprehensive dataset was then split into an 80% training set and a 20% testing set for model development and evaluation, respectively. The XGBoost classifier was configured with the following key parameters: a multiclass classification objective, the number of target classes set to the total number of peptides, 1000 boosting rounds (trees), and a learning rate of 0.05 to control the step size shrinkage. Regularization was applied with max depth of 5 to limit tree complexity, minimum child weight was 1 to control minimum sum of instance weight (hessian) needed in a child, gamma was set to 0 for minimum loss reduction required to make a further partition on a leaf node, subsample was 0.8 (fraction of samples used per tree), and the fraction of features used per tree was 0.8. L1 and L2 regularization were used. For reproducibility, a random state was fixed, and computation was distributed across all available CPU cores. The model’s performance during training was monitored using the multiclass classification error. The trained model’s performance was evaluated on the unseen testing dataset. Classification metrics including accuracy, precision, recall, and F1-score were summarized in a standard classification report, and a confusion matrix was generated to visualize per-class prediction accuracy.

### Multistage ensemble probabilistic blending classifier

A multistage ensemble probabilistic blending classifier was developed to identify guest-host peptides based on translocation events through a set of nanopore variants. This hierarchical model architecture was designed to leverage the distinct information provided by different nanopore mutations **(Fig. 4A)**. All classifiers within the ensemble were implemented using the XGBoost algorithm. The model was trained in three distinct stages: a set of Stage 1 binary classifiers, a set of Stage 2 peptide-specific classifiers, and a final meta-classifier. The overall dataset, comprising translocation features from multiple nanopore variants (WT, F427A, and F427Y), was first subjected to a single, global train-test split with an 80:20 ratio. This split was performed with stratification to ensure that the distribution of both peptide and nanopore classes was maintained proportionally in both the training and testing sets.

#### Stage 1 classifiers: aromatic/non-aromatic discrimination

The first stage of the model consisted of training a separate binary classifier for each nanopore variant used. Each of these Stage 1 classifiers was a nanopore-specialist model, trained exclusively on data from its respective nanopore. Their objective was to perform a coarse-grained classification of the guest-host peptides into two broad categories: aromatic (e.g., Phe, Trp, TrpDL, Tyr) and non-aromatic (e.g., Ala, Leu, Thr). The probabilistic outputs from these Level 1 classifiers, specifically the probability of a translocation event belonging to the ‘aromatic’ class, were used as new features for the subsequent stages.

#### Stage 2 classifiers: peptide-specific classification

The outputs from the Level 1 classifiers were then used to augment the original feature set of the entire training data. The augmented dataset, now including the Level 1 probabilities, was used to train two separate Level 2 classifiers. The data was partitioned based on the Level 1 prediction: one Level 2 classifier was trained on all data points predicted as ‘aromatic’ and was tasked with classifying the specific aromatic peptide, while the other was trained on data predicted as ‘non-aromatic’ to classify the specific non-aromatic peptide. These classifiers refined the coarse Stage 1 predictions into fine-grained, per-peptide class predictions.

#### Meta-classifier: probabilistic blending

The final stage of the ensemble was a meta-classifier. This model was trained on a blended feature set consisting of the probabilistic outputs from both the Stage 1 and Stage 2 classifiers. For each translocation event, the meta-classifier’s input features included the aromatic/non-aromatic probabilities from each nanopore’s Stage 1 classifier, as well as the peptide-specific probabilities from the Stage 2 classifiers. By training on these probabilistic outputs, the meta-classifier learned to weigh the predictions of the preceding stages, effectively blending their outputs to produce a final, robust prediction for the specific guest-host peptide class. Classification metrics including accuracy, precision, recall, and F1-score were summarized in a standard classification report, and a confusion matrix was generated to visualize per-class prediction accuracy.

## Acknowledgments

We thank members of the department for useful feedback and discussions. J.M.C. and B.A.K. conceived of the experiments. J.M.C. collected the data. B.A.K performed the analysis. B.A.K. and J.M.C. wrote the manuscript. Portions of this document, including some of the Python code and language refinement, were generated with the assistance of AI-powered tools. All content was reviewed and approved by the authors, who take full responsibility for its accuracy.

---

**Table S1.** Performance of single-stage XGBoost classification model for individual nanopores^1^.

| Nanopore | Overall Accuracy<sup>2</sup> | Macro-averaged F1-score<sup>2</sup> | F1-score per guest-host peptide class<sup>3</sup> |  |  |  |  |  |  |
| --- | --- | --- | --- | --- | --- | --- | --- | --- | --- |
|  |  |  | Ala | Leu | Phe | Thr | Trp | TrpDL | Tyr |
| WT | 0.8659<br>(±0.0087) | 0.8652<br>(±0.0087) | 0.80 | 0.91 | 0.93 | 0.74 | 0.93 | 0.96 | 0.87 |
| F427A | 0.8874<br>(±0.0089) | 0.8869<br>(±0.0089) | 0.95 | 0.82 | 0.82 | 0.94 | 0.96 | 0.91 | 0.87 |
| F427Y | 0.7856<br>(±0.0085) | 0.7829<br>(±0.0088) | 0.93 | 0.73 | 0.60 | 0.89 | 0.80 | 0.88 | 0.70 |

<sup>1</sup>Performance based on test set evaluation metrics of single-stage XGBoost classification model.
<sup>2</sup>Overall accuracy and macro-averaged F1-scores are means and std. dev. (N=5).
<sup>3</sup>Best set of per-class F1-scores out of N=5 replicates.

**Fig. S1.**
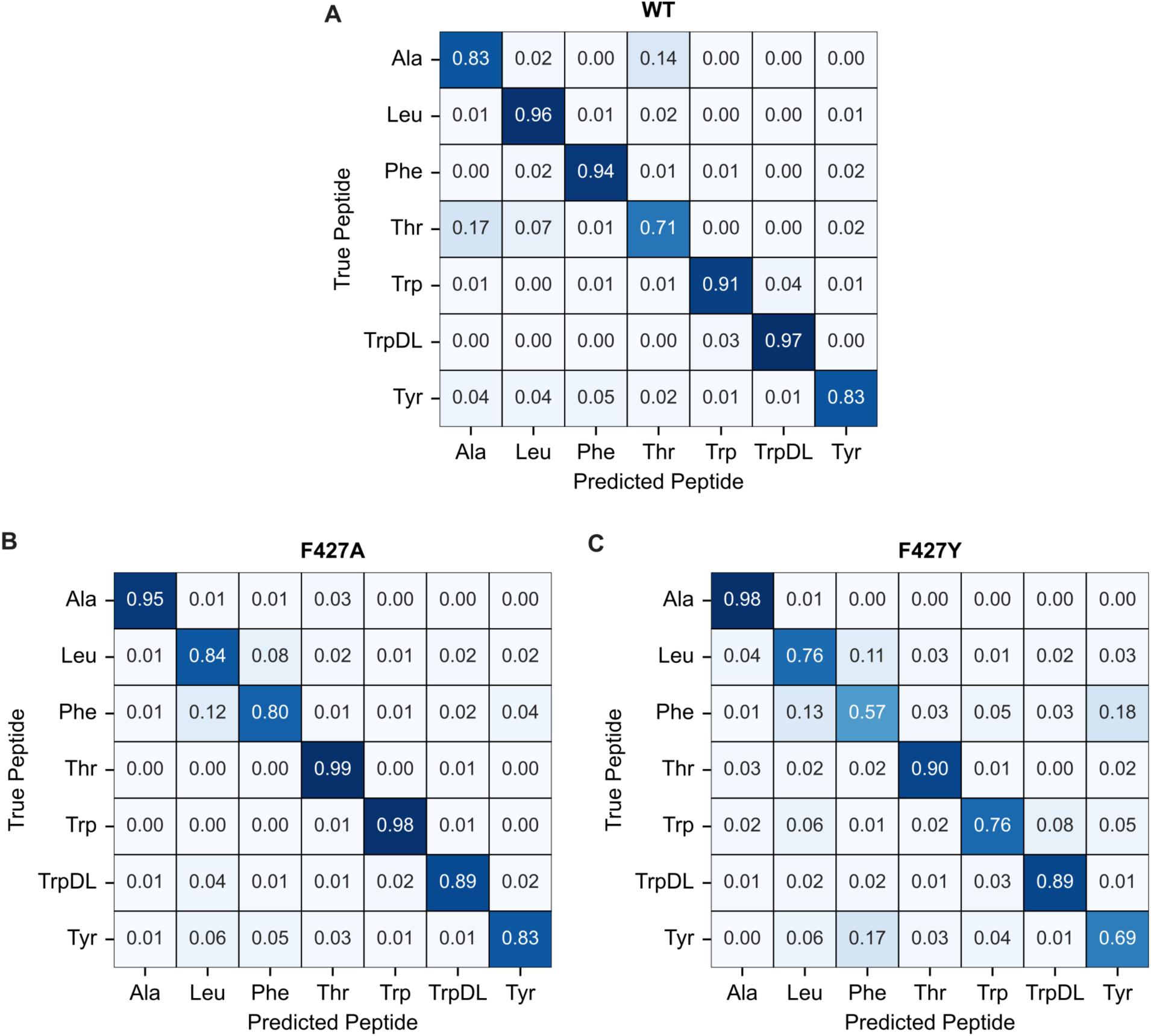
Confusion matrices for nanopore ϕ-clamp variants using a single-stage XGBoost classification. Normalized confusion matrices from most successful translocation event classifications using a single-stage XGBoost model for **(A)** WT, **(B)** F427A, and **(C)** F427Y nanopores. Rows represent true peptide labels, and columns represent predicted peptide labels. Values indicate the proportion of events from a given true class that were predicted as each class.

